# Genomic and physiological analyses reveal that extremely thermophilic *Caldicellulosiruptor changbaiensis* deploys unique cellulose attachment mechanisms

**DOI:** 10.1101/622977

**Authors:** Asma M.A.M. Khan, Carl Mendoza, Valerie J. Hauk, Sara E. Blumer-Schuette

**Affiliations:** Department of Biological Sciences, Oakland University, Rochester, MI 48309

**Keywords:** *Caldicellulosiruptor*, extreme thermophile, cellulase, tāpirin

## Abstract

The genus *Caldicellulosiruptor* are extremely thermophilic, heterotrophic anaerobes that degrade plant biomass using modular, multifunctional enzymes. Prior pangenome analyses determined that this genus is genetically diverse, with the current pangenome remaining open, meaning that new genes are expected with each additional genome sequence added. Given the high biodiversity observed among the genus *Caldicellulosiruptor*, we have sequenced and added a 14^th^ species, *Caldicellulosiruptor changbaiensis*, to the pangenome. The pangenome now includes 3,791 ortholog clusters, 120 of which are unique to *C. changbaiensis* and may be involved in plant biomass degradation. Comparisons between *C. changbaiensis* and *Caldicellulosiruptor bescii* on the basis of growth kinetics, cellulose solubilization and cell attachment to polysaccharides highlighted physiological differences between the two species which are supported by their respective gene inventories. Most significantly, these comparisons indicated that *C. changbaiensis* possesses unique cellulose attachment mechanisms not observed among the other strongly cellulolytic members of the genus *Caldicellulosiruptor*.

## INTRODUCTION

The genus *Caldicellulosiruptor* is comprised of extremely thermophilic, fermentative heterotrophs whose members have been isolated worldwide from terrestrial geothermal springs or other thermal environments [37]. The original isolates from the genus *Caldicellulosiruptor* were identified on the basis of their ability to grow on cellulose at elevated temperatures [56,54], especially temperatures beyond the optimal growth temperature of *Ruminiclostridium thermocellum* [48]. Interest in thermostable enzymes produced by this genus continues, as the initial discovery of their multifunctional, modular enzymes [51,26,57,67] represented an alternate paradigm to cellulosomes [2,52]. Further discoveries on the capabilities of these thermostable enzymes include the unique mode of action used by the central cellulase, CelA, [8], synergistic activity in ionic liquid optimized enzyme mixtures [45,46] and the creation of designer cellulosomes from *Caldicellulosiruptor* catalytic domains [29]. Development of a genetics system for *Caldicellulosiruptor bescii* [14,16] has also expanded the scope of work with this genus, including metabolic engineering [10,12,13,50] and catalytic improvement [18,30,32,31,34,33].

The availability of genome sequences has precipitated deeper insights into the genus *Caldicellulosiruptor*, including comparative studies which have identified biomarkers for plant biomass deconstruction [6,5,23], novel insertion elements [15], genetic tractability [11], diverse mechanisms involved in biomass solubilization [66,37], unique cellulose adhesins (tāpirins) [5,37] and the identification of new combinations of catalytic domains [5,36,23]. Perhaps owing to the unique thermal environments that this genus inhabits, their genomes appear to be dynamic, as the first described *Caldicellulosiruptor* pangenome was predicted to be open [5], and remained open after the addition of five additional genome sequences [36].

Here, we have analyzed the genome sequence of *Caldicellulosiruptor changbaiensis*, isolated from a hot spring in the Changbai Mountains [3], representing the 14^th^ and most recent addition to the *Caldicellulosiruptor* pangenome. Past *Caldicellulosiruptor* pangenomes were comprised of multiple species from most countries of origin, which allowed for prior analysis on the basis of biogeography [5], with the exception of China and Japan [20]. Now with the addition of the *C. changbaiensis* genome sequence, insights into the biogeography of isolates from China and how they compare to the global *Caldicellulosiruptor* pangenome is possible. Furthermore, on the basis of the open *Caldicellulosiruptor* pangenome [20,5], we hypothesize that the *C. changbaiensis* genome may encode for novel substrate-binding proteins and/ or plant biomass degrading enzymes. In addition to updating the *Caldicellulosiruptor* pangenome, we also present differences in the growth physiology of *C. changbaiensis* versus *Caldicellulosiruptor bescii*, currently the benchmark species against which most *Caldicellulosiruptor* are compared for their plant biomass degrading capabilities.

## MATERIALS AND METHODS

### Microbial strains and medium

Freeze-dried stocks of *C. changbaiensis* strain CBS-Z were obtained from the Leibniz Institute DSMZ – German Collection of Microorganisms and Cell Cultures (DSMZ). Glycerol stocks of *C. bescii* DSM-6725 were obtained from the laboratory of Robert M. Kelly, North Carolina State University (Raleigh, NC). Both species were cultured at 75°C on low osmolarity defined (LOD) medium [25] under a nitrogen headspace to maintain anaerobic conditions and supplemented with carbohydrates as a carbon source. Carbohydrates used as a carbon source included cellobiose (≥ 99%, Chem-Impex Int’l, Inc.), pectin (Sigma-Aldrich), xylan (Sigma-Aldrich), glucomannan (NOW Foods), and microcrystalline cellulose (20 µm Sigmacell, Sigma-Aldrich). For genomic DNA isolation, *C. changbaiensis* was cultured anaerobically at 75°C on low osmolarity complex (LOC) medium [25] with cellobiose as a carbon source.

### Genomic DNA isolation

Genomic DNA was isolated using the Joint Genome Institute’s CTAB-based protocol (https://jgi.doe.gov/user-programs/pmo-overview/protocols-sample-preparation-information/jgi-bacterial-dna-isolation-ctab-protocol-2012/), with modifications. In order to isolate enough DNA for sequencing, 500 ml of overnight *C. changbaiensis* culture was harvested by centrifugation at 5000x*g*, 4°C for 20 minutes and resuspending the cell pellet in 14.8 ml of TE buffer, prior to lysis. Gel electrophoresis in 0.7% agarose was used to assess the quality of genomic DNA and the concentration and purity of the sample for sequencing was quantified using a NanoDrop spectrophotometer, and Qubit fluorometric assay (dsDNA HS assay, Thermo Fisher). Prior to genome sequencing, a 16S rRNA gene fragment was amplified from isolated genomic DNA using oligonucleotide primers (Eton Bioscience) previously designed for identification of *C. changbaiensis* [3], for positive identification of *C. changbaiensis* (Table 1). Amplicons were sent for Sanger sequencing (Eton Bioscience), using the same oligonucleotide primers.

**Table 1.**
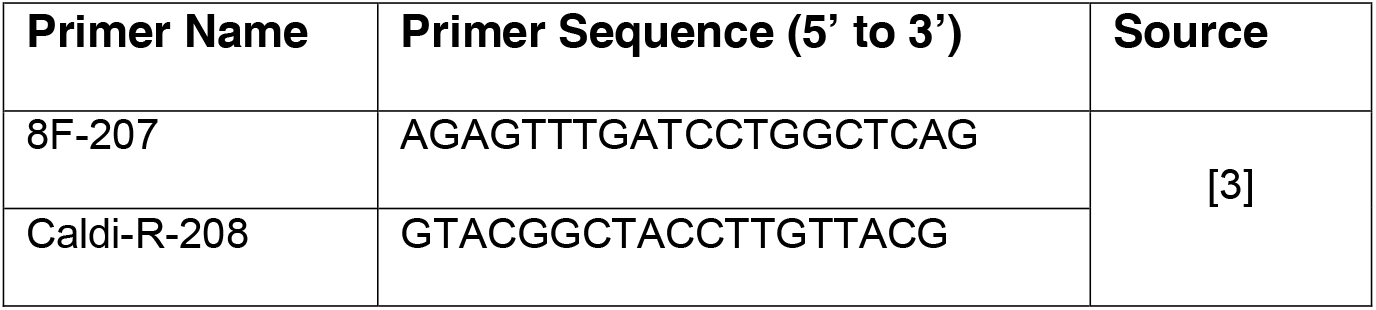
Oligonucleotide primers used for *Caldicellulosiruptor* 16S rRNA gene fragment amplification.

### *C. changbaiensis* genome sequencing, assembly and annotation

The genome sequence for *C. changbaiensis* [40] was assembled to 60-fold coverage from long-read Oxford NanoPore (MinION) data generated in house, and short-read Illumina data generated by Molecular Research, LP (MR DNA). Hybrid assembly of the complete *C. changbaiensis* genome used Unicycler v0.4.7 [61], and annotation of the genome used the Prokaryotic Genome Annotation Pipeline v4.7 [55] provided by the National Center for Biotechnology Information (NCBI). The assembled genome and reads used for assembly of the *C. changbaiensis* genome are available through NCBI BioProject accession PRJNA511150.

### Phylogenomic analysis

Fourteen genome sequences from the genus *Caldicellulosiruptor* were included in the phylogenomic analyses (see Table 2 for genome sequence accession numbers). Orthologous protein groups were classified using the GET_HOMOLOGUES v20092018 software package [19], running OrthoMCL v1.4 [39], COGtriangles v2.1 [35], or bidirectional best hits (BDBH) as determined by BLASTP [1,9]. Orthologous protein clusters were determined using the OrthoMCL parameters: 75% pairwise coverage, maximum BLASTP E-value of 1e-5, and MCL inflation of 1.5. GET_HOMOLOGUES was also used to parse the pangenome matrices comparing the *C. changbaiensis* genome inventory against the recent 13 *Caldicellulosiruptor* pangenome [37] or the revised *C. bescii* genome [22]. Core-(Eq. 1) and pangenome (Eq. 2) parameters were predicted after curve fitting randomly sampled core- or pangenome data to functions previously described by Tettelin et al., [58].

**Table 2.** *Caldicellulosiruptor* genome sequences included in the updated pangenome analysis.

| **Species Name** | **NCBI RefSeq Accession** | **Reference** |
| --- | --- | --- |
| *C. acetigenus* | GCF_000421725.1 | [41] |
| *C. bescii* | GCF_000022325.1 | [22] |
| *C. changbaiensis* | GCF_003999255.1 | [40] |
| *C. danielii* | GCF_000955725.1 | [38,36] |
| *C. hydrothermalis* | GCF_000166355.1 | [7,5] |
| *C. kristjanssonii* | GCF_000166695.1 | [7,5] |
| *C. kronotskyensis* | GCF_000166775.1 | [7,5] |
| *C. lactoaceticus* | GCF_000193435.2 | [7,5] |
| *C. morganii* | GCF_000955745.1 | [38,36] |
| *C. naganoensis* | GCF_000955735.1 | [38,36] |
| *C. obsidiansis* | GCF_000145215.1 | [24] |
| *C. owensensis* | GCF_000166335.1 | [7,5] |
| *C. saccharolyticus* | GCF_000016545.1 | [59] |
| *C. str. F32* | GCF_000404025.1 | [63] |

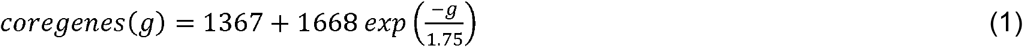

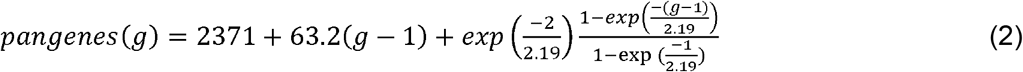

Genome-level similarity was quantified as average nucleotide identity (ANIb) from the BLASTN+ alignment of 1,020 nt fragments from the 14 *Caldicellulosiruptor* genomes [49,27]. ANIb were calculated by Pyani v.0.2.7, (https://github.com/widdowquinn/pyani) and percent identities were plotted as a heatmap by the software package.

### Growth kinetics on polysaccharides

*C. bescii* or *C. changbaiensis* were revived from −80°C glycerol stocks for growth curve analysis on microcrystalline cellulose, xylan, pectin or glucomannan. Glycerol stocks (1 ml) were subcultured into 50 ml LOD medium for 3 consecutive subcultures using 2% (v/v) inoculum at each passage. Revived cultures were then transferred (2% [v/v] inoculum) to LOD medium containing a 1:1 ratio of maltose (*C. bescii*) or cellobiose (*C. changbaiensis*) to polysaccharide. The 1:1 mixture was then passaged (2% [v/v] inoculum) three times successively in LOD medium with polysaccharide, only. Cultures for growth curves were inoculated at a starting cell density of 1 × 10^6^ cells ml^−1^ in 200 ml LOD plus the respective polysaccharide. Biological replicates were used for each growth phase experiment. Cell counting used epifluorescence microscopy at 1000x total magnification and a counting reticle as described previously [28]. Cells were fixed in a final volume of 1.1 ml gluteraldehyde (2.5% [v/v] in water) prior to incubation with acridine orange (1 g l^−1^) and approximately 5 ml sterilized water and thoroughly mixed. Stained cells were then vacuum filtered through a polycarbonate 0.22 µm filter (GE). Samples were counted using a 10×10 reticle a total of ten times. Cell counts were averaged for calculation of cell density (cells ml^−1^). Doubling times are described as the number of hours per generation during exponential growth, calculated as Δtime divided by the number of generations.

### Microcrystalline cellulose solubilization

Solubilization of microcrystalline cellulose followed protocols established by Zurawski *et al*., [66] with modifications. *C. bescii* or *C. changbaiensis* were cultured in serum bottles with 50 ml of LOD medium supplemented with 0.6g of microcrystalline cellulose (20µm Sigmacell) at a starting cell density of 10^6^ cells ml^−1^. Cultures were then incubated without shaking at 75°C for seven days, after which the remaining microcrystalline cellulose was harvested by centrifugation at 6000 xg, 4°C for 15 min in a swing bucket rotor. The cellulose pellet was washed four times in sterile, deionized water and air dried at 75°C until the weight of the microcrystalline cellulose did not change. Uninoculated LOD served as an abiotic control. Percent solubilization is reported as the difference in substrate weight divided by the starting weight multiplied by 100. All experimental conditions were measured in triplicate and significance was determined by a t-test (p-value < 0.05).

### Cell attachment assays

*C. bescii* and *C. changbaiensis* cell cultures were grown to early stationary phase on either xylan or cellulose (1 g l^−1^) as the carbon source, and cell densities were calculated before harvesting at 5000 xg for 10 minutes at room temperature. Cells were resuspended and concentrated ten-fold in the binding buffer (50 mM sodium phosphate, pH 7.2) to a 10-fold density of approximately 1-2 × 10^9^ cells ml^−1^ for cells cultured on xylan or 1 × 10^8^ cells mil^−1^ for cells cultured on cellulose. For each treatment condition, 1.2 ml of *C. bescii* or *C. changbaiensis* planktonic cells in binding buffer were added to a 1.5 ml microcentrifuge tube, and supplemented with 10 mg of washed substrate (experimental condition: xylan or cellulose), or no substrate for the negative control. All assay tubes were incubated at room temperature for one hour with gentle rotary shaking at 100 rpm. After incubation, planktonic cells were enumerated as described above for the growth curves. Each binding assay was repeated six times. Two-sample t-tests were used to analyze the data using the R studio statistics package v.3.3.3 [47].

## RESULTS AND DISCUSSION

### Phylogenomic analysis of the *C. changbaiensis* genome

With the addition of the fourteenth *Caldicellulosiruptor* genome [40], we sought to define an updated core- and pangenome. Three different algorithms: OrthoMCL [39], bidirectional best hit and COGtriangles [35] were used to classify orthologous clusters for pangenome analysis (**Table S1**). Of the three, the clusters formed by OrthoMCL resulted in an estimated core- and pangenome with the lowest residual standard errors, and are reported here (Fig. 1). Overall, there are 120 unique protein clusters identified in the *C. changbaiensis* genome when compared to the prior *Caldicellulosiruptor* pangenome [37], 75 of which were annotated as hypothetical proteins. Further transcriptomic and proteomic studies may aid in the identification of the function of these unique hypothetical proteins. By adding a 14^th^ genome, the *Caldicellulosiruptor* core genome was reduced to 1,367 orthologous clusters (see Eq. 1), however, the pangenome (3,791 genes) continues to expand at an estimated rate of 63.2 genes per additional genome (Eq. 2, Fig. 1) highlighting the plasticity of the *Caldicellulosiruptor* pangenome.

**Figure 1.**
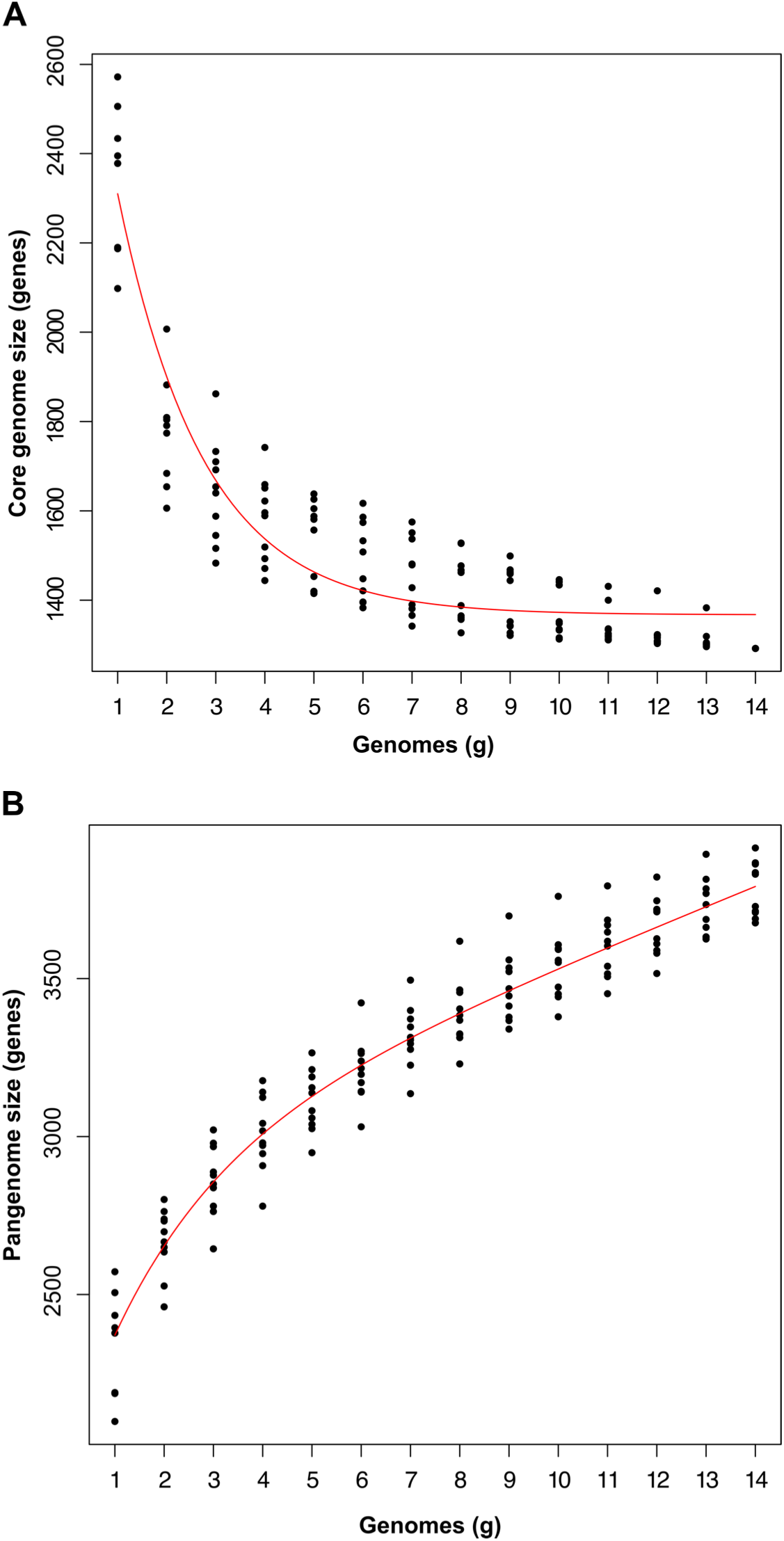
Core- and pangenome size estimates calculated from random sampling of 14 *Caldicellulosiruptor* genomes. **(a)** Fitted curve of the estimated *Caldicellulosiruptor* core genome from 10 random samples of genomes up to n=14. The current size of the core genome is 1367 orthologous clusters. **(b)** Fitted curve of the estimated *Caldicellulosiruptor* pangenome from 10 random samples of genomes up to n=14. The *Caldicellulosiruptor* pangenome remains open and has increased to 3791 genes. The rate of growth for the pangenome is 63.2 new genes per genome sequenced. Core- and pangenome estimates were calculated from the equations reported by Tettelin *et al*., [58] using GET_HOMOLOGUES software [19].

In contrast to previously released genome sequences from New Zealand [36], *C. changbaiensis* exhibits a similar pattern of biogeography based on average nucleotide identity (ANIb). As expected, *Caldicellulosiruptor* sp. F32, isolated from compost in China [63], and *C. naganoensis*, isolated from a hot spring in Japan [56] shared higher percent identity levels with *C. changbaiensis*, along with *C. saccharolyticus*, isolated from a hot spring in New Zealand (Fig. 2, **Table S2**). All species that *C. changbaiensis* shared the highest ANI with have been described and confirmed as being strongly cellulolytic, implying that the *C. changbaiensis* genome would also encode for a glucan degradation locus (GDL). Despite the high level of ANIb, based on the open *Caldicellulosiruptor* pangenome, we expected to find new genes involved in carbohydrate metabolism and possibly GDL arrangements.

**Figure 2.**
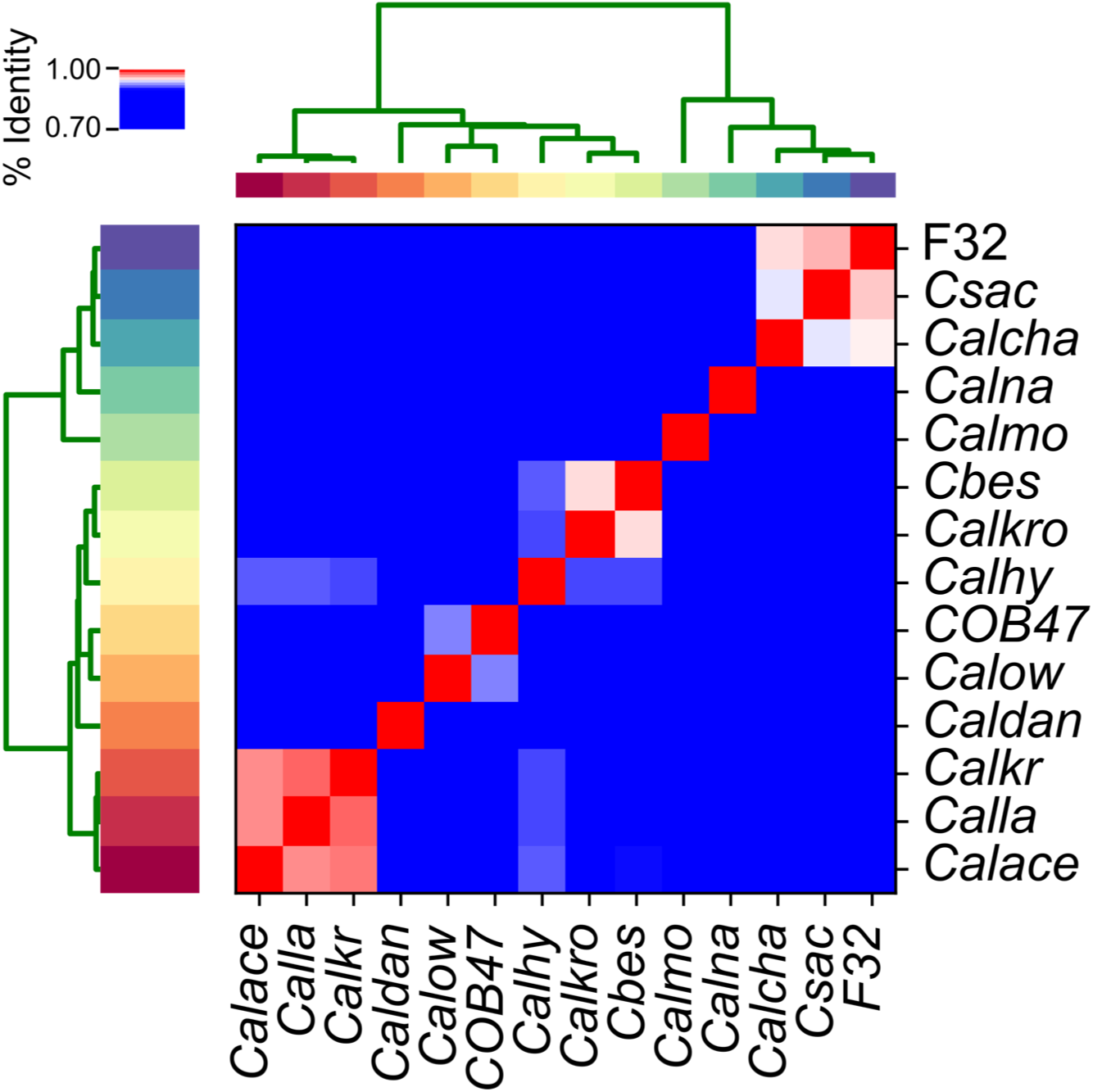
Heatmap representation of the average nucleotide identity for 14 genome sequenced species from the genus *Caldicellulosiruptor*. Average nucleotide identity (ANIb) was calculated on the basis of legacy BLASTn sequence identity over 1020nt sequence fragments. ANIb values of all 14 genomes are represented by a heat plot ranging from blue (75%< ANIb <90%), white (90%< ANIb <95%) to red (ANIb >95%). Pyani (https://github.com/widdowquinn/pyani) was used to calculate ANIb values and generate the clustered heatmap. Hierarchal cluster dendrograms were generated on the basis of similar ANIb values across each species. ANIb values are reported in **Table S1**. Calace, *C. acetigenus*; Cbes, *C. bescii*; Calcha, *C. changbaiensis*; Caldan, *C. danielii*; Calhy, *C. hydrothermalis*; Calkr, *C. kristjanssonii*; Calkro, *C. kronotskyensis*; Calla, *C. lactoaceticus*; Calmo, *C. morganii*; Calna, *C. naganoensis*; COB47, *C. obsidiansis*; Calow, *C. owensensis*; Csac, *C. saccharolyticus*; F32, *C*. sp. F32.

### *C. changbaiensis* exhibits different abilities to grow on polysaccharides versus *C. bescii*

In order to benchmark the ability of *C. changbaiensis* to grow on plant-related polysaccharides, we compared its doubling time during exponential growth on representative plant polysaccharides to *C. bescii* (Table 3). Doubling times (generation time) were calculated from cell densities measured during exponential growth. Overall, *C. changbaiensis* grows slower on microcrystalline cellulose than *C. bescii*, with a 38% larger doubling time during growth on crystalline cellulose, however, both cultures grew at similar rates on xylan. On both glucomannan, and pectin, *C. changbaiensis* grew faster with 35% lower doubling times (Table 3). The differential ability of *C. changbaiensis* and *C. bescii* to grow on pectin and glucomannan is not unexpected, as the differential ability from one species to another to hydrolyze and metabolize plant biomass, comprised of polysaccharides such as xylan, pectin and glucomannan, was previously observed, in one case *C. saccharolyticus* grew slower on plant biomass versus *C. bescii* [62] and *C. kronotskyensis* [66] and another observation where *C. danielii* grew approximately 50% faster than *C. bescii*, *C. morganii* and *C. naganoensis* on plant biomass [36].

**Table 3.**
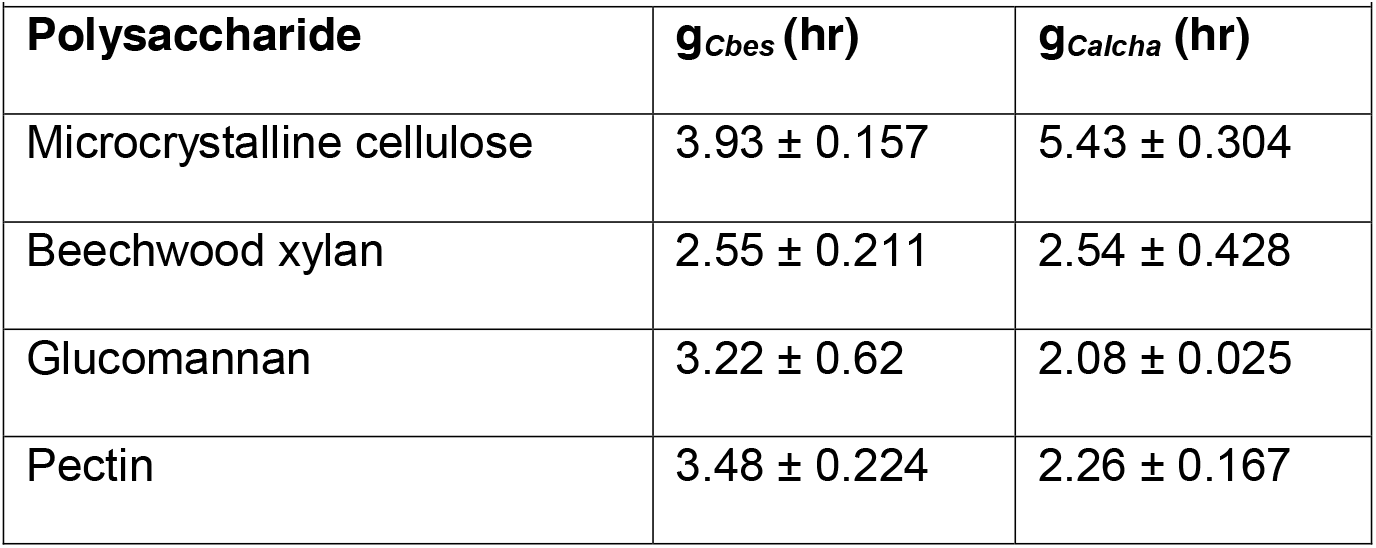
Doubling time of *C. changbaiensis* or *C. bescii* grown on plant polysaccharides.

|  |  |  |
| --- | --- | --- |
| Polysaccharide | $g_{Cbes}$ (hr) | $g_{Calcha}$ (hr) |
| Microcrystalline cellulose | $3.93 \pm 0.157$ | $5.43 \pm 0.304$ |
| Beechwood xylan | $2.55 \pm 0.211$ | $2.54 \pm 0.428$ |
| Glucomannan | $3.22 \pm 0.62$ | $2.08 \pm 0.025$ |
| Pectin | $3.48 \pm 0.224$ | $2.26 \pm 0.167$ |

When comparing the genomes of *C. changbaiensis* and *C. bescii*, *C. changbaiensis* encodes for 411 genes not shared with *C. bescii*, 120 of which are unique to the genus. We expect that the differences in growth rates on carbohydrates to be related to differences in gene inventory. In fact, the *C. changbaiensis* gene inventory encoding for carbohydrate active enzymes includes 13 genes not found in the *C. bescii* genome, including an annotated β-mannanase (glycoside hydrolase [GH] family 26) and two mannooligosaccharide phosphorylases (GH130). This additional β-mannanase and phosphorylases likely contribute to the enhanced growth of *C. changbaiensis* on glucomannan (Table 3).

The lower doubling time on pectin is surprising, however, given that *C. changbaiensis* does not encode for the pectinase cluster that is located in the *C. bescii* genome immediately downstream of the GDL. *C. bescii* gene deletion strains lacking the pectinase cluster were impaired in their growth on both pectin-rich plant biomass and pectin [17], indicating that *C. changbaiensis* has evolved alternate mechanisms to deconstruct or metabolize pectin. Screening the *C. changbaiensis* genome for pectin-related enzymes did not identify any genes encoding for polysaccharide lyases (PL) that were unique in comparison to *C. bescii*, however genes encoding for representatives from GH family 43, 51 (α-L-arabinofuranosidases) and 95 (α-fucosidase) were present. One scenario is that these enzymes participate in the hydrolysis of carbohydrate sidechains from pectin [44]. Another plausible explanation is that *C. changbaiensis* has evolved to import and efficiently ferment a broader range of carbohydrates released during growth on plant biomass, including uronic acids, and/ or the deoxy sugars fucose and rhamnose. While *C. bescii* may rely on its enzymatic repertoire to deconstruct plant biomass, it may not metabolize all types of carbohydrates that are released, similar to *R. thermocellum* which produces xylanases, but does not metabolize xylose [42,43].

### Organization of the *C. changbaiensis* genome degradation locus

*C. changbaiensis* was originally described as strongly cellulolytic [3] and accordingly, its genome encodes for a GDL that shares a similar organization with other strongly cellulolytic members of the genus. since *C. bescii* was able to grow at a faster rate on microcrystalline cellulose than *C. changbaiensis* (Table 3), we opted to focus on the comparison of GDL between these two species. The GDL from both species is remarkedly similar, with only CelD possessing a different arrangement of catalytic and non-catalytic domains (GH10-CBM3-GH5) from *C. changbaiensis*, and truncated versions of CelE (GH9-CBM3-GH5) and CelF (GH74-CBM3) present (Fig. 3). Prior *in vitro* biochemical analyses on the synergy of cellulase mixtures from *C. bescii* had observed that a mixture of three cellulases, CelA, CelC and CelE (ACE cellulases) worked synergistically to hydrolyze cellulose as well as a mixture of all six *C. bescii* cellulases [21]. One could hypothesize, then, that members of the genus *Caldicellulosiruptor* that possess all three of these enzymes would be among the most cellulolytic. Three additional species, *C. kronotskyensis*, *C. danielii*, and *C. naganoensis* also share a similar organization of their GDL [36], including the presence of CelA, CelC and CelE. The contributions of CelD and CelF to cellulose hydrolysis or solubilization are low [22,21] and likely not to impact the ability of *C. changbaiensis* to efficiently hydrolyze cellulose.

**Figure 3.**
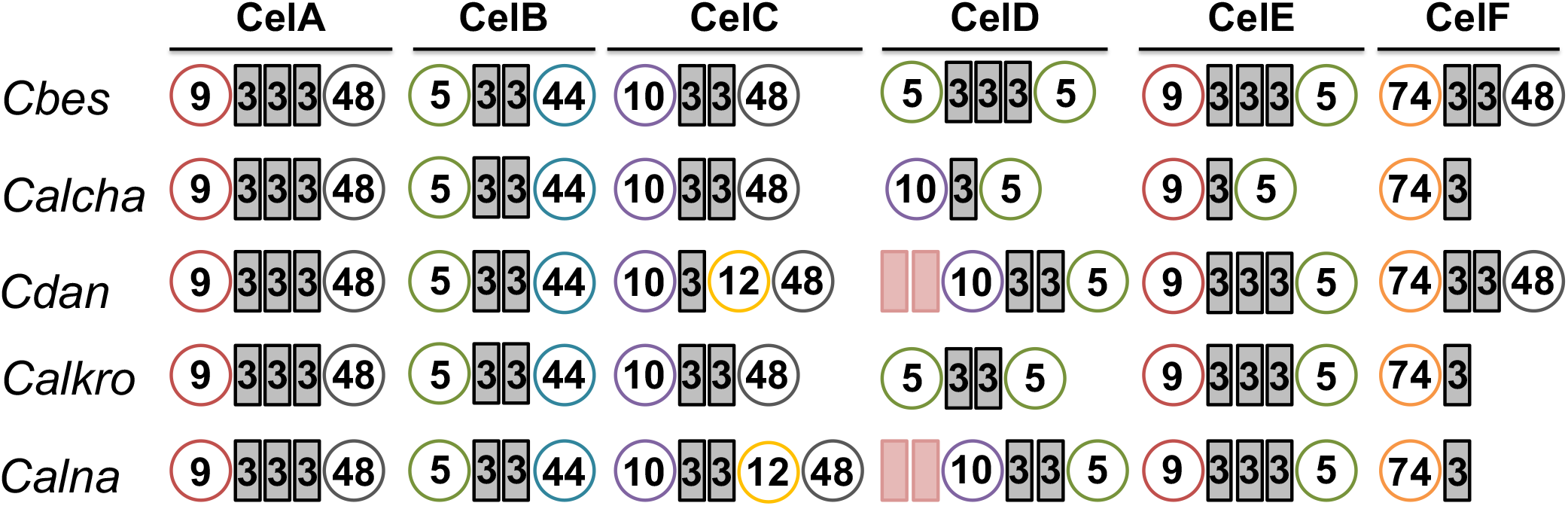
Modular multifunctional enzymes encoded for by the glucan degradation locus. Glucan degradation loci were selected on the basis of the presence of “ACE” cellulases. ACE cellulases: CelA, CelC and CelE. Circles represent the glycoside hydrolase (GH) domains, rectangles represent the carbohydrate binding module (CBM) domains. GH5, green circles; GH9, red circles; GH10, violet circles; GH 44, blue circles; GH48, grey circles; GH74, orange circles. CBM3, grey rectangles; CBM22, pink rectangles.

Indeed, *C. changbaiensis* can solubilize microcrystalline cellulose (Fig. 4), however the amount of cellulose solubilized was 22.4% lower than the amount solubilized by *C. bescii*, which is similar to the performance of *C. saccharolyticus* when compared to *C. bescii* [24.8% lower, 66]. This result begs the question if the mere presence of the ACE cellulases is sufficient to meet the *C. bescii* benchmark for hydrolysis of cellulose. One explanation could be that the *C. changbaiensis* CelE ortholog may not be as efficient in cellulose hydrolysis since it is lacking two CBM3 modules. However, the nearly equal reduction of cellulose solubilization by both *C. bescii* gene deletion strains incapable of producing CelA-CelC versus CelA-CelE does not support this possibility [22]. Furthermore, CelE truncations that possessed the GH9 catalytic domain and three or two CBM3 domains were equally capable of microcrystalline cellulose hydrolysis [53], making it unlikely that the loss of a CBM3 domain from the *C. changbaiensis* CelA ortholog hampered its activity.

**Figure 4.**
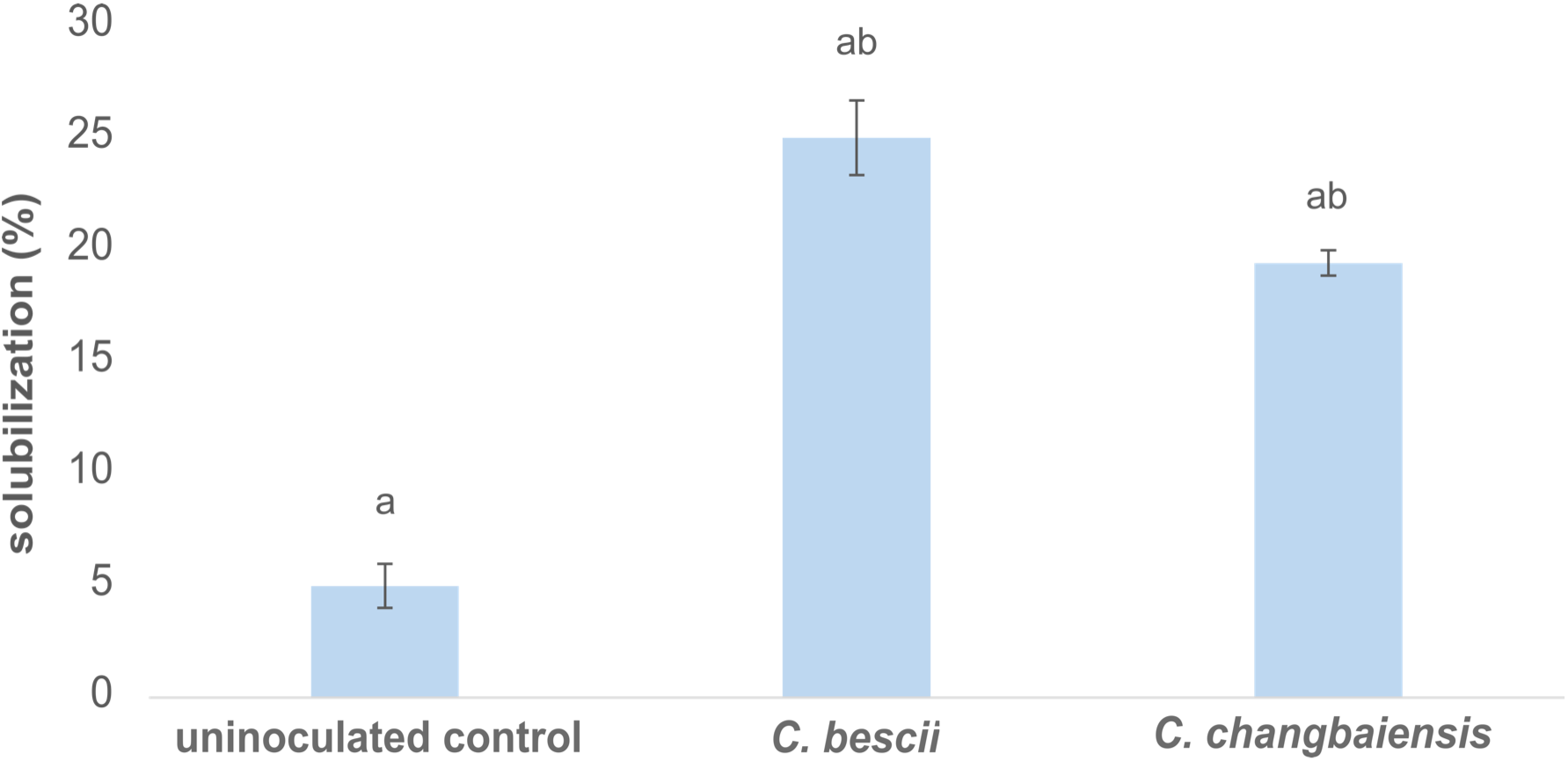
Solubilization of microcrystalline cellulose by *C. bescii* and *C. changbaiensis*. Uninoculated control, indicates abiotic cellulose solubilization in LOD medium. Error bars represent standard error (n=3). Similar letters over columns denote p< 0.05 as determined by a t-test.

Alternately, sequence divergence of ACE cellulase orthologs may play a larger role in the catalytic capacity of cellulolytic members from the genus *Caldicellulosiruptor*. Of the ACE cellulases, CelA is a key player, supported by its unique hydrolysis mechanism [8], the severe reduction in cellulose hydrolysis by *C. bescii* celA gene deletion mutant [65,22], and biochemical analysis of GDL enzyme synergy [21]. Prior comparison of CelA orthologs from *C. bescii* and *C. danielii* found *Cb*CelA to be a superior enzyme [36], indicating that GDL sequences have diverged during speciation, making it likely that the ACE cellulases from *C. changbaiensis* may not demonstrate the same catalytic efficiency as *C. bescii*.

### Attachment of *C. bescii* and *C. changbaiensis* to plant polysaccharides

Aside from comparisons of catalytic ability, we also compared the ability of *C. changbaiensis* versus *C. bescii* planktonic cells to bind to insoluble substrates (xylan and cellulose). A decrease in the planktonic cell density (PCD) after exposure to the substrate compared to the PCD of the negative controls without substrate is indicative of cells binding to the substrate. Surprisingly, we saw no such decrease in PCD for *C. changbaiensis* cultured on xylan after incubation with cellulose or xylan (Figs. 5A and B). This inability of *C. changbaiensis* to attach to xylan or cellulose after growth on xylan is surprising, given that xylan is a major polysaccharide constituent of lignocellulose, and would likely serve as a chemical signal. Since no xylan or cellulose attachment proteins are produced in response to growth on xylan, *C. changbaiensis* appears to act as a specialist, responding only to cellulose. Regardless, when *C. changbaiensis* is grown on cellulose, it maintains an ability to attach to cellulose (29% cells attached), which is slightly lower than the relative amount of *C. bescii* cells attached to cellulose (33% attached, Fig. 5C). Surprisingly, when *C. bescii* cells cultured on xylan were tested for attachment to either xylan or cellulose there was a significant decrease in (PCD) of indicating that *C. bescii* cells grown on xylan are producing proteins capable of attaching to xylan (33% attachment, Fig. 5A) or cellulose (68% attachment, Fig. 5B). While we expected to see cells from cultures grown on xylan attaching to xylan, interestingly, *C. bescii* cell attachment was most pronounced when cells were grown on xylan and incubated with cellulose (Fig. 5B). The ability of *C. bescii* to attach to cellulose (Figs. 5B and C), is in large part due to the presence of tāpirins, since a *C. bescii* tāpirin deletion mutant was severely impaired in cellular attachment to cellulose [37].

**Figure 5.**
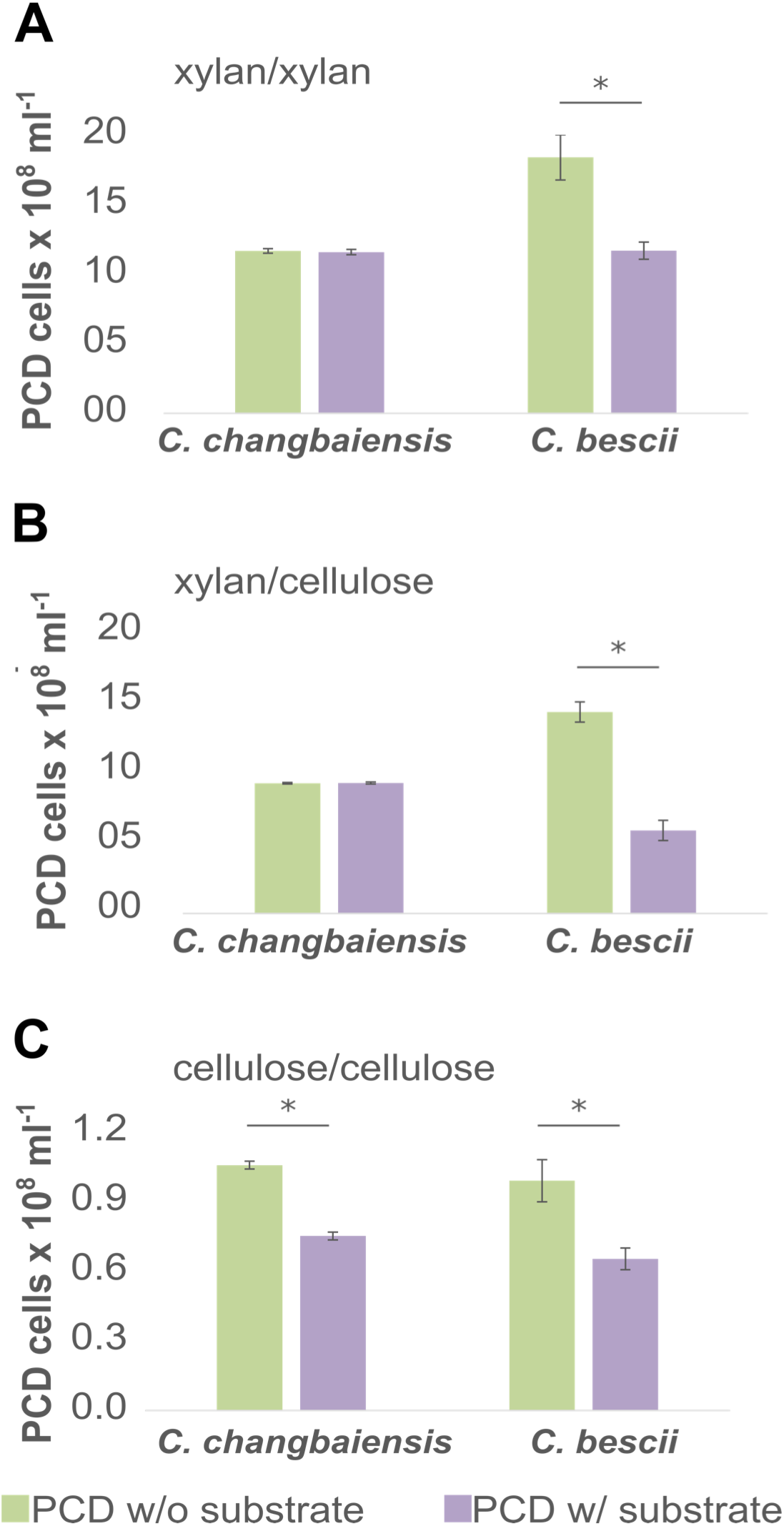
Comparison of the ability of *C. bescii* or *C. changbaiensis* planktonic cells to attach to polysaccharides. Titles above bar charts indicate the carbon source for growth/ binding substrate. **(a, b)** When cells are grown on xylan, only planktonic *C. bescii* cells were able to attach to xylan or cellulose. **(c)** Cells grown on cellulose as the carbon source and exposed to cellulose as the binding substrate. Planktonic cell densities (PCD), enumerated by epifluorescence microscopy are plotted on the y-axis. Green columns indicate PCD without binding substrate and purple columns indicate PCD with the binding substrate. * indicates p < 0.01 as determined by a t-test. All assays had n=6 biological replicates.

### The *C. changbaiensis* genome encodes for atypical tāpirin genes

Another notable difference observed between *C. changbaiensis* and *C. bescii* during growth on cellulose is the lack of floc formation by *C. changbaiensi*s (Fig. 6). Based on this discrepancy between *C. changbaiensis* and *C. bescii*, we examined the genomic context of the type IV pilus locus encoded by the *C. changbaiensis* genome (Fig. 7). The T4P locus is found in the genome in all members of the *Caldicellulosiruptor*, and is also located upstream of the GDL in the genomes of strongly cellulolytic species [5,4]. Most notably, while a full T4P locus is present in the *C. changbaiensis* genome, classical tāpirin genes are absent which encode for proteins that bind with high affinity to cellulose [4,37]. Instead, two genes with little, to no homology to the classical tāpirins are located directly downstream of the T4P locus which we will refer to as atypical tāpirins. The proteins encoded for by these genes are not unique to *C. changbaiensis*, as both *C. acetigenus* and *C. ownesensis* also encode for these atypical tāpirins. All three species encode for two atypical tāpirins: a hypothetical protein (Genbank accession: WP_127352232.1) and a von Willebrand Factor A protein (Genbank accession: WP_127352233.1) (yellow arrows, Fig. 7). While *C. changbaiensis* shares a similar genomic context at the 3’ end of the T4P locus, the atypical *C. changbaiensis* tāpirins are not close orthologs, as they share 74.33% and 68.01% amino sequence similarity with the first and second atypical tāpirins encoded *C. owensensis*. Prior proteomics data collected from cellulose-bound, supernatant and whole cell lysate protein fractions determined that both atypical tāpirins are produced by *C. owensensis* in response to cellulose [5], supporting their potential role in cell attachment to cellulose.

**Figure 6.**
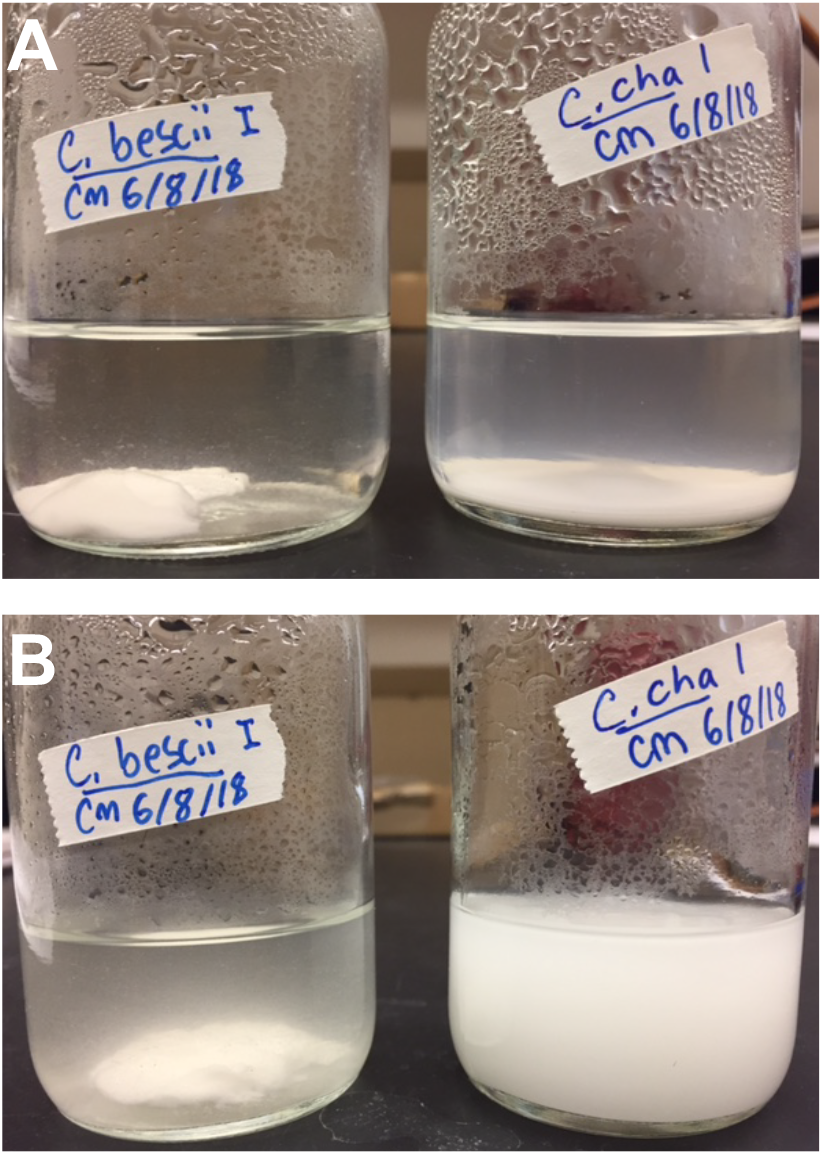
Flocculation of *C. bescii* cells cultured on chemically defined medium and microcrystalline cellulose. **(a)** Formation of a floc of *C. bescii* cells around microcrystalline cellulose (diameter, 20µm) while planktonic *C. changbaiensis* cells (cloudiness) are visible. **(b)** Same serum bottles as in “A”, however the bottles were vigorously mixed. The *C. bescii* floc remains fairly stable, while both microcrystalline cellulose and cells are mixed in the *C. changbaiensis* culture.

**Figure 7.**
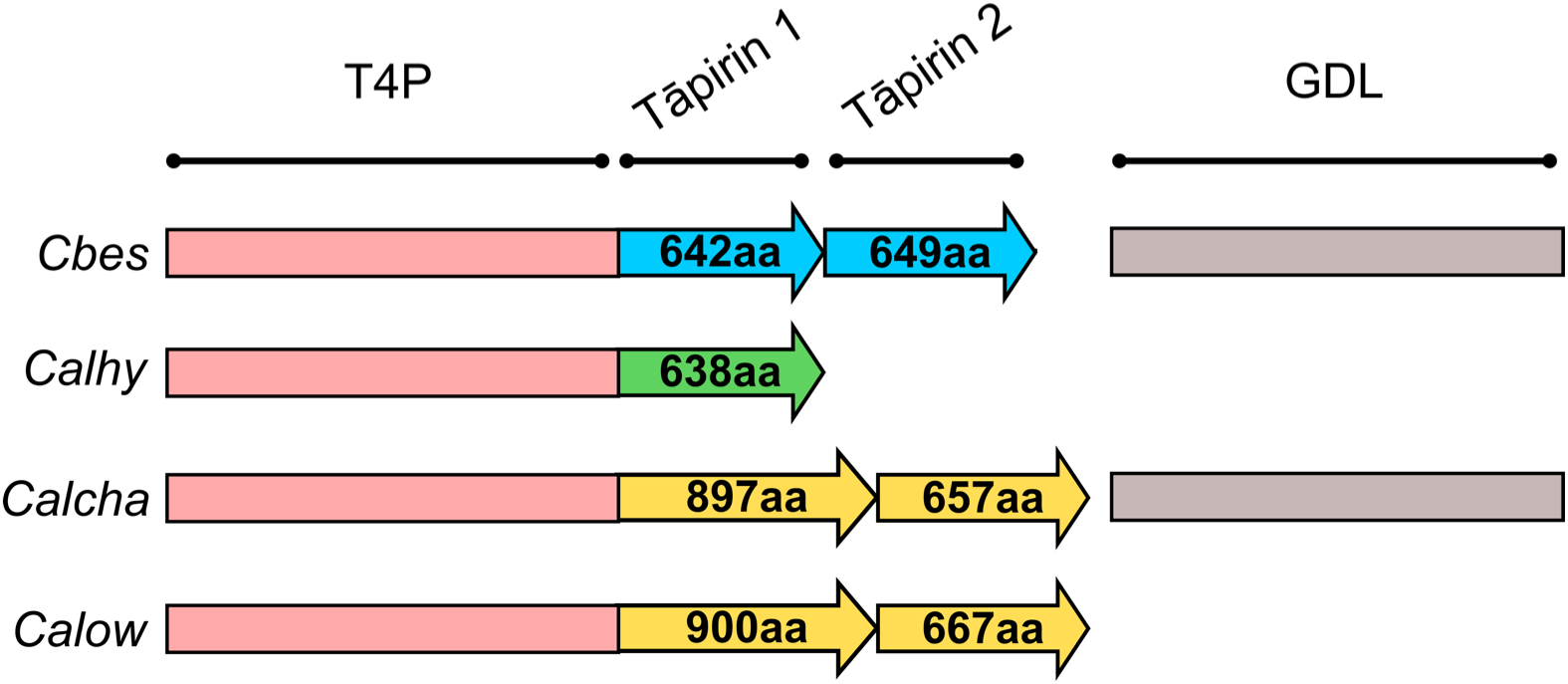
Genomic context for the location of the tāpirins from strongly to weakly cellulolytic *Caldicellulosiruptor* species. Different colors represent the classical versus atypical tāpirins. Blue arrows: Cbes tāpirin 1 (Gen bank accession: YP_002573732) and Cbes tāpirin 2 (Gen bank accession: YP_002573731). Green arrow: Calhy tāpirin 1 (Gen bank accession number: YP_003992006). Yellow arrows: Calcha tāpirin 1 (Gen bank accession: WP_127352232.1) and 2 (Gen bank accession: WP_127352233.1), and Calow tāpirin 1 (Gen bank accession: YP_004002936) and 2 (Gen bank accession r: YP_004002935). Grey rectangles indicate the presence of the GDL downstream of the tāpirins. Atypical tāpirin 1 is annotated as a hypothetical protein and atypical tāpirin 2 is annotated as a von Willebrand factor A protein. Cbes, *C. bescii*; Calhy, *C. hydrothermalis*; Calcha, *C. changbaiensis* and Calow, *C. owensensis*. Peach rectangles represent the type IV pilus locus directly upstream of the tāpirins. Arrows indicate tāpirin 1 and 2. Numbers in the tāpirin arrows indicate the amino acid length.

This observed sequence divergence between the atypical tāpirins from strongly and weakly cellulolytic species is similar to the tāpirin encoded for by *C. hydrothermalis* which shares little amino acid sequence homology with classical tāpirins, but shares a similar tertiary structure, and is capable of occupying more sites on crystalline cellulose in comparison to classical tāpirins [37]. Production of tāpirins with an affinity to cellulose likely plays a role in the ability of weakly cellulolytic members of the genus to adhere to cellulose and benefit from the cellooligosaccharides released by the action of cellulases [60]. The atypical tāpirins, originally only observed in the genomes of weakly cellulolytic species, may also serve as cellulose adhesins, however, further in-depth biochemical characterization of both atypical tāpirin proteins is required to confirm their function.

## CONCLUSIONS

Overall, the *Caldicellulosiruptor* pangenome remains open, and is expected to gain approximately 63 new genes with each additional species sequenced (Fig. 1A). The addition of a second species isolated from China indicates that the diversity of *Caldicellulosiruptor* species from this region is higher than those isolated from Iceland, however, the level of observed diversity is not as high as those species isolated from Kamchatka, Russia or New Zealand on the basis of ANIb (Fig. 2). *C. changbaiensis* encodes for a GDL (Fig. 3) similar in organization as *C. bescii*, however is not as cellulolytic as *C. bescii* on the basis of doubling time (Table 3) and cellulose solubilization (Fig. 4). However, *C. changbaiensis* does appear to have a broader metabolic appetite for uronic acids or deoxy sugars. *C. changbaiensis* also fails to form a floc during growth on microcrystalline cellulose (Fig. 6), a phenotype previously described for C. bescii [64], however both species are capable of attaching to cellulose (Fig. 5). Interestingly, *C. bescii* retains an ability to attach to cellulose when previously grown on xylan, while *C. changbaiensis* does not (Fig. 5B) indicating that the two species respond differently to soluble carbohydrates present in their environment. Tāpirins were previously demonstrated to be key cellulose adhesins for strongly [4] to weakly cellulolytic [37] members of the genus *Caldicellulosiruptor*. Surprisingly, *C. changbaiensis* does not encode for the classical tāpirins, and instead encodes for atypical tāpirins, one of which possesses a von Willebrand type A protein domain (Fig. 7). These atypical tāpirins are homologous to those encoded for by weakly cellulolytic *C. owensensis* and *C. acetigenus*, however this may not indicate that the atypical tāpirins are not involved in attachment to cellulose, as the divergent classical tāpirin encoded for by *C. hyrothermalis* binds at a high density to cellulose [37]. The combined lack of classical tāpirins, along with the ability to attach to cellulose indicates that *C. changbaiensis* evolved a unique strategy to attach to cellulose. Further study on the biophysical properties of these atypical tāpirins is warranted to assess their ability to interact with plant polysaccharides, including cellulose.

## Acknowledgements

The authors wish to acknowledge the support of an Oakland University Provost Award, awarded to C. Mendoza.

