## Supplementary Data for "Genomic and physiological analyses reveal that extremely thermophilic *Caldicellulosiruptor changbaiensis* deploys unique cellulose attachment mechanisms"

---

Asma M.A.M. Khan‡, Carl Mendoza‡, Valerie J. Hauk and Sara E. Blumer-Schuette*

**Journal:** J Ind Microl Biotechnol

*Corresponding author: Sara E. Blumer-Schuette
Dept. of Biological Sciences
Oakland University


**Table S1. Comparison of the number of ortholog clusters identified by three different algorithms**

**Table S2. Average nucleotide identity values calculated from legacy BLASTn**

**Table S1. Comparison of the number of ortholog clusters identified by three different algorithms**

| | Core-genome (genes)[b] | SE[a] | Pangenome (genes) | SE |
|---|---|---|---|---|
| OrthoMCL[c] | 1367.8 | 99.49 | 3791.1 | 104.2 |
| COG | 1380.1 | 100.41 | 3779.6 | 106.28 |
| BDBH | 1363.5 | 101.36 | 3833.4 | 103.95 |

[a] Residual standard error

[b] Core- and pangenome size estimates from curve fitting after Tettelin *et al*. [4]

[c] OrthoMCL [3], COG [2] and BDBH algorithms were run using GET_HOMOLOGUES [1].

**Table S2. Average nucleotide identity values calculated from legacy BLASTn**

| | Calow | Calmor | Calkro | Calla | Csac | Calcha | Caldan | F32 | Calhy | Calace | Calkr | Calna | Cbes | COB47 | |
|---|---|---|---|---|---|---|---|---|---|---|---|---|---|---|---|
| **Calow** | 1.000 | 0.762 | 0.872 | 0.886 | 0.777 | 0.778 | 0.863 | 0.776 | 0.893 | 0.885 | 0.885 | 0.774 | 0.876 | 0.925 | *C. owensensis* OL |
| **Calmor** | 0.763 | 1.000 | 0.769 | 0.775 | 0.824 | 0.823 | 0.795 | 0.822 | 0.770 | 0.767 | 0.769 | 0.830 | 0.775 | 0.770 | *C. morganii* Rt8.B8 |
| **Calkro** | 0.871 | 0.764 | 1.000 | 0.894 | 0.795 | 0.795 | 0.870 | 0.794 | 0.913 | 0.896 | 0.894 | 0.790 | 0.955 | 0.878 | *C. kronotskyensis* 2002 |
| **Calla** | 0.882 | 0.770 | 0.891 | 1.000 | 0.787 | 0.792 | 0.877 | 0.784 | 0.913 | 0.970 | 0.980 | 0.783 | 0.895 | 0.890 | *C. lactoaceticus* 6A |
| **Csac** | 0.776 | 0.824 | 0.795 | 0.790 | 1.000 | 0.942 | 0.805 | 0.959 | 0.795 | 0.784 | 0.787 | 0.880 | 0.803 | 0.785 | *C. saccharolyticus* DSM 8903 |
| **Calcha** | 0.777 | 0.821 | 0.798 | 0.790 | 0.944 | 1.000 | 0.807 | 0.952 | 0.799 | 0.788 | 0.786 | 0.884 | 0.805 | 0.789 | *C. changbaiensis* CBS-Z |
| **Caldan** | 0.859 | 0.792 | 0.870 | 0.880 | 0.805 | 0.808 | 1.000 | 0.807 | 0.888 | 0.884 | 0.878 | 0.797 | 0.874 | 0.864 | *C. danielii* Wai35.B1 |
| **F32** | 0.774 | 0.820 | 0.794 | 0.781 | 0.962 | 0.953 | 0.808 | 1.000 | 0.796 | 0.782 | 0.782 | 0.888 | 0.798 | 0.781 | *C.* sp. F32 |
| **Calhy** | 0.890 | 0.770 | 0.914 | 0.917 | 0.792 | 0.798 | 0.884 | 0.796 | 1.000 | 0.915 | 0.913 | 0.782 | 0.913 | 0.896 | *C. hydrothermalis* 108 |
| **Calace** | 0.884 | 0.765 | 0.897 | 0.971 | 0.784 | 0.790 | 0.885 | 0.781 | 0.918 | 1.000 | 0.973 | 0.777 | 0.900 | 0.889 | *C. acetigenus* DSM 7040 |
| **Calkr** | 0.881 | 0.765 | 0.893 | 0.979 | 0.785 | 0.788 | 0.875 | 0.782 | 0.911 | 0.969 | 1.000 | 0.781 | 0.893 | 0.886 | *C. kristjanssonii* I77R1B |
| **Calna** | 0.771 | 0.825 | 0.794 | 0.787 | 0.884 | 0.886 | 0.793 | 0.888 | 0.782 | 0.782 | 0.787 | 1.000 | 0.796 | 0.781 | *C. naganoensis* NA10 |
| **Cbes** | 0.875 | 0.770 | 0.955 | 0.898 | 0.803 | 0.805 | 0.874 | 0.799 | 0.914 | 0.897 | 0.894 | 0.795 | 1.000 | 0.880 | *C. bescii* DSMZ 6725 |
| **COB47** | 0.923 | 0.764 | 0.879 | 0.891 | 0.784 | 0.787 | 0.864 | 0.782 | 0.896 | 0.888 | 0.888 | 0.780 | 0.881 | 1.000 | *C. obsidiansis* OB47 |
| | **Calow** | **Calmor** | **Calkro** | **Calla** | **Csac** | **Calcha** | **Caldan** | **F32** | **Calhy** | **Calace** | **Calkr** | **Calna** | **Cbes** | **COB47** | |
